# Prediction of Eye, Hair and Skin Color in Admixed Populations of Latin America

**DOI:** 10.1101/2020.12.09.415901

**Authors:** Sagnik Palmal, Kaustubh Adhikari, Javier Mendoza-Revilla, Macarena Fuentes-Guajardo, Caio C. Silva de Cerqueira, Juan Camilo Chacón-Duque, Anood Sohail, Malena Hurtado, Valeria Villegas, Vanessa Granja, Claudia Jaramillo, William Arias, Rodrigo Barquera Lozano, Paola Everardo-Martínez, Jorge Gómez-Valdés, Hugo Villamil-Ramírez, Tábita Hünemeier, Virginia Ramallo, Rolando Gonzalez-José, Lavinia Schüler-Faccini, Maria-Cátira Bortolini, Victor Acuña-Alonzo, Samuel Canizales-Quinteros, Carla Gallo, Giovanni Poletti, Gabriel Bedoya, Francisco Rothhammer, David Balding, Pierre Faux, Andrés Ruiz-Linares

## Abstract

We report an evaluation of prediction accuracy for eye, hair and skin pigmentation based on genomic and phenotypic data for over 6,500 admixed Latin Americans (the CANDELA dataset). We examined the impact on prediction accuracy of three main factors: (i) The methods of prediction, including classical statistical methods and machine learning approaches, (ii) The inclusion of non-genetic predictors, continental genetic ancestry and pigmentation SNPs in the prediction models, and (iii) Compared two sets of pigmentation SNPs: the commonly-used HIrisPlex-S set (developed in Europeans) and novel SNP sets we defined here based on genome-wide association results in the CANDELA sample. We find that Random Forest or regression are globally the best performing methods. Although continental genetic ancestry has substantial power for prediction of pigmentation in Latin Americans, the inclusion of pigmentation SNPs increases prediction accuracy considerably, particularly for skin color. For hair and eye color, HIrisPlex-S has a similar performance to the CANDELA-specific prediction SNP sets. However, for skin pigmentation the performance of HIrisPlex-S is markedly lower than the SNP set defined here, including predictions in an independent dataset of Native American data. These results reflect the relatively high variation in hair and eye color among Europeans for whom HIrisPlex-S was developed, whereas their variation in skin pigmentation is comparatively lower. Furthermore, we show that the dataset used in the training of prediction models strongly impacts on the portability of these models across Europeans and Native Americans.

## Introduction

There is growing interest in the use of genetic data for the prediction of physical appearance, particularly in forensic, historical and paleo-anthropological studies[1–3]. Strong impetus for these studies has been provided by Genome Wide Association Studies (GWAS) of traits such as pigmentation, which have enabled the identification of single nucleotide polymorphisms (SNPs) associated with eye, hair and skin color[4–6]. Early prediction studies evaluated the accuracy of predicting eye color from SNP data[7–10] and these analyses have been more recently extended to hair [11,12], and skin pigmentation[13]. As a result, sets of SNPs have now been proposed for the simultaneous prediction of eye, hair and skin color[14,15]. However, so far, these studies have had a strong European bias[7,10,11,14,16]. The pigmentation SNPs for prediction have been selected from GWAS performed mainly in Northern Europeans[4,5,17] and prediction models also optimized mostly in European datasets[17–21]. Furthermore, evaluation of the accuracy of these prediction tools has also been performed essentially in European-derived population samples[18–20,22].

Among major continental populations, Europeans show the largest variation in eye and hair color, whereas the variation in skin color in Europeans is relatively lower compared to other continental regions such as Africa or South Asia[23–28]. The other continental regions have narrower variation in eye and hair color, mostly within the brown-black range; although fine-scale quantitative measurement still allows us to discern this variability and identify genetic variants unique to those populations[27,29–31]. Interestingly, even though the underlying biological mechanisms are broadly similar for these different types of pigmentations, their genetic architecture is substantially different, as eye and hair pigmentation are chiefly controlled by a handful of large-effect variants, whereas skin pigmentation is observed to be more polygenic and the impact of any single variant is smaller[6,16,24–27]. Consequently, studies assessing the portability of pigmentation prediction systems derived primarily on European cohorts have reported good portability for eye and hair phenotypes in other non-European continental populations, where most samples were correctly predicted to be in the brown-black category, whereas portability for skin color in other continental populations was poorer[23,32,33]; for example, all light-skinned Japanese individuals were incorrectly predicted to be of dark skin color[23]. The convergent evolution of lighter skin color in different regions of Eurasia, as reported previously in the CANDELA cohort[31] and elsewhere[26,34,35], is the likely cause for this discrepancy, and highlights the importance of using diverse cohorts for building skin pigmentation prediction models.

Latin Americans represent one of the largest recently admixed populations worldwide. The history of Latin America has involved extensive admixture, particularly between Native Americans, Europeans and sub-Saharan Africans (with respective estimates of ancestry share: 38%, 52% and 6%[36]). Consistent with its partly European ancestry, a recent GWAS for pigmentation traits in Latin Americans in the CANDELA cohort detected phenotypic effects for a number of loci previously identified in Europeans[31]. In addition to these, novel pigmentation SNPs with genome-wide significant association were also identified in that study. These included SNPs polymorphic only in East Asians and Native Americans, consistent with the independent evolution of skin pigmentation in West and East Eurasia[26,34,35]. The admixed ancestry of Latin America and the finding in the region of pigmentation variants not present in Europeans emphasizes the need to evaluate the accuracy of tools currently available for prediction of pigmentation traits in this population.

Here we aimed to evaluate the accuracy of prediction of pigmentation traits in Latin Americans in the large Latin American dataset (the CANDELA cohort) recently used in GWASs of physical appearance, including for hair, eye and skin pigmentation[31,36–41]. We focused on comparing the accuracy of prediction methods, the predictors included in models, and the SNP sets used for prediction. We find that Random Forest and regression are generally the method with best performance, depending on the predicted trait. We also find that the inclusion of pigmentation SNPs increases the accuracy of prediction models substantially over that obtained by the inclusion of other predictors. Finally, the HIrisPlex-S has a similar performance to a CANDELA-specific set of prediction SNPs except for skin pigmentation.

## Materials and Methods

### Study sample: phenotypes, genetic data and covariates

We analyzed data previously studied by the CANDELA consortium for GWAS of pigmentation traits[31,36–39]. The consortium gathered genetic and phenotypic data from over 6,500 individuals recruited in five Latin American countries: Mexico (N=~1,200), Colombia (N=~1,700), Peru(N=~1,230), Chile (N=~1,730) and Brazil (N=~630).

Pigmentation traits evaluated directly on the research subjects consists of (A) hair pigmentation (recorded in four categories: 1-red/reddish, 2-blond, 3-dark blond/light brown or 4-brown/black. However, due to their very low frequency (<0.6%), individuals in the ‘red/reddish’ category were not included here), (B) eye color, recorded as five ordered categories: 1-blue/gray, 2-honey, 3-green, 4-light brown, 5-dark brown/black. For increasing consistency with previous publications[10,13,42–44], here we recoded these data into just three categories: 1-Blue/Gray, 2-Intermediate (honey or green) and 3-Brown/Black (light brown or dark brown/black) and (C) a quantitative measure of skin pigmentation from an area unexposed to sunlight (the Melanin Index MI, obtained by reflectometry). We also had available additional measures of iris pigmentation, extracted from digital photographs, using the HCL color space (Hue, Chroma and Luminance). Hue being an angle (recorded in arc degree), we linearized this trait with cosine and shifted the angle by 15° in order to maximize the number of samples in the range [0,180°]; hence the trait considered is cos(Hue+15). The frequency distribution for these traits in the CANDELA dataset is shown in Supplementary Figure S1.

For comparison with other studies[13,42], we converted the skin MI into a three-level categorical trait. For this, we selected CANDELA individuals with 100% European ancestry (N=70) and examined their facial photographs in order to label them as very pale or pale. Similarly, we selected CANDELA individuals with highest African ancestry (>39%; N=23) and examined their photographs in order to label them as dark or very dark. We plotted the distribution of the MI for individuals in the very pale, pale, dark and very dark categories and identified MI values of 33 and 47 as thresholds for categorical skin color: fair (MI <33; N= 2,506), intermediate (MI 33-47; N=3,840) and dark (MI>47; N=180) classified as dark (see Supplementary figure S2). These thresholds are in line with values obtained in another study of admixed Brazilians[45].

The genetic data consisted of ~9 million genotypes, ~700k of which were obtained experimentally by genotyping Illumina’s Omni Express chip, the remainder obtained by imputation as described in Adhikari et al[31]. We applied several filters to the CANDELA dataset prior to the trait prediction analyses. Firstly, we retained only individuals aged 18 to 45. Secondly, we removed 8 pairs of individuals whose pairwise probability of IBD was estimated close to 1, to discard potential sample mix-ups (hence, 16 individuals removed), and individuals whose estimated African ancestry was more than European and native ancestry estimations (23 individuals), as those were considered as genetic outliers and thus excluded. Finally, we excluded all individuals with missing data on any of the covariates (age, sex, BMI). Note that BMI was considered as a covariate since we found it significantly correlated to some pigmentation phenotypes; that correlation is most likely a confounding effect of continental genetic ancestry. The final sample size used in the analyses was: 6,495 for hair color, 6,526 for MI, 6,529 for categorical eye color and 5,738 for quantitative eye color traits – Hue, Chroma and Luminance. These three eye color phenotypes constitute the bicone color space model – HCL scale for human perception of eye colors (previously explained by Adhikari et al[31]).

### Pigmentation SNP sets used for prediction

We used two sets of SNPs for the prediction analyses of pigmentation traits.

Firstly, as a benchmark we used HIrisplex-S, a SNP set that has been developing over the years for the prediction of eye, hair and skin pigmentation mainly in Europeans[7,11,14] HIrisplex-S currently includes 41 SNPs, 22 of which were directly genotyped in the CANDELA samples (genotypes for the remaining 19 SNPs being imputed). In our analyses, we only retained SNPs with MAF >= 1% in the CANDELA data, reducing the HIrisplex-S set used here to 34 SNPs (see Supplementary table S1).

Secondly, we devised “CANDELA” (CAN) SNP sets for prediction of each pigmentation trait (E-eye; H-hair; S-skin) based on results from a GWAS conducted in the CANDELA sample[31]. To pre-select SNPs for each trait, we used the following protocol: (1) selection of all SNPs with GWAS association *p*-values <1E-5, (2) grouping (clumping) of SNPs in high LD and (3) for each SNP group, selection of the SNP with highest predictive power (it is well-known that the most significant marker is not necessarily the best predictive marker[46]). In this way we pre-selected 1,471 SNPs relevant for skin pigmentation prediction, 207 for hair pigmentation prediction and 701 for eye pigmentation prediction. These SNPs were then ranked based on conditional predictive power for each trait, in order to search for the set of predictive SNPs with the least SNPs included. Details of the approach used are provided in Supplementary method S1, and the resulting CAN-E, CAN-S and CAN-H sets described in the Results section.

### Prediction methods and models evaluated

A broad array of statistical methods have been employed in the literature to predict pigmentation traits, such as (multiple) linear[17,47] or (multinomial) logistic regression[10,11], decision trees[17,48], neural networks[17,49], and naïve Bayes classifiers[33,42,50]. Each method has its advantages and disadvantages, and are better suited for certain types of traits, e.g. linear regression for quantitative traits[17,47] and logistic regression for categorical traits[10,11].

The overall strategy we used for performing pigmentation prediction in the CANDELA dataset is shown in Figure 1. Linear Regression (LR) or Multinomial Logistic Regression (MLR) were used as the reference methods for quantitative or categorical traits, respectively. These two methods were used to evaluate three prediction models, incorporating an increasing number of predictors (Fig. 1A):

1. Using only non-genetic covariates as predictors:

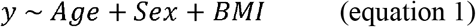
2. Incorporating genetic ancestry to model 1:

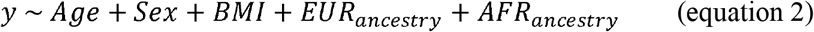 Here we included as predictors the estimates of European and African ancestry (obtained by unsupervised admixture estimation on genome-wide data). Native American ancestry was omitted so as to avoid collinearity (since the three continental ancestries sum to 1).
3. Incorporating pigmentation SNPs to model 2:

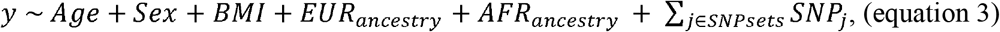

where SNPsets refers to SNPs included either in the HIrisPlex-S or CAN sets defined above.

**Figure 1.**
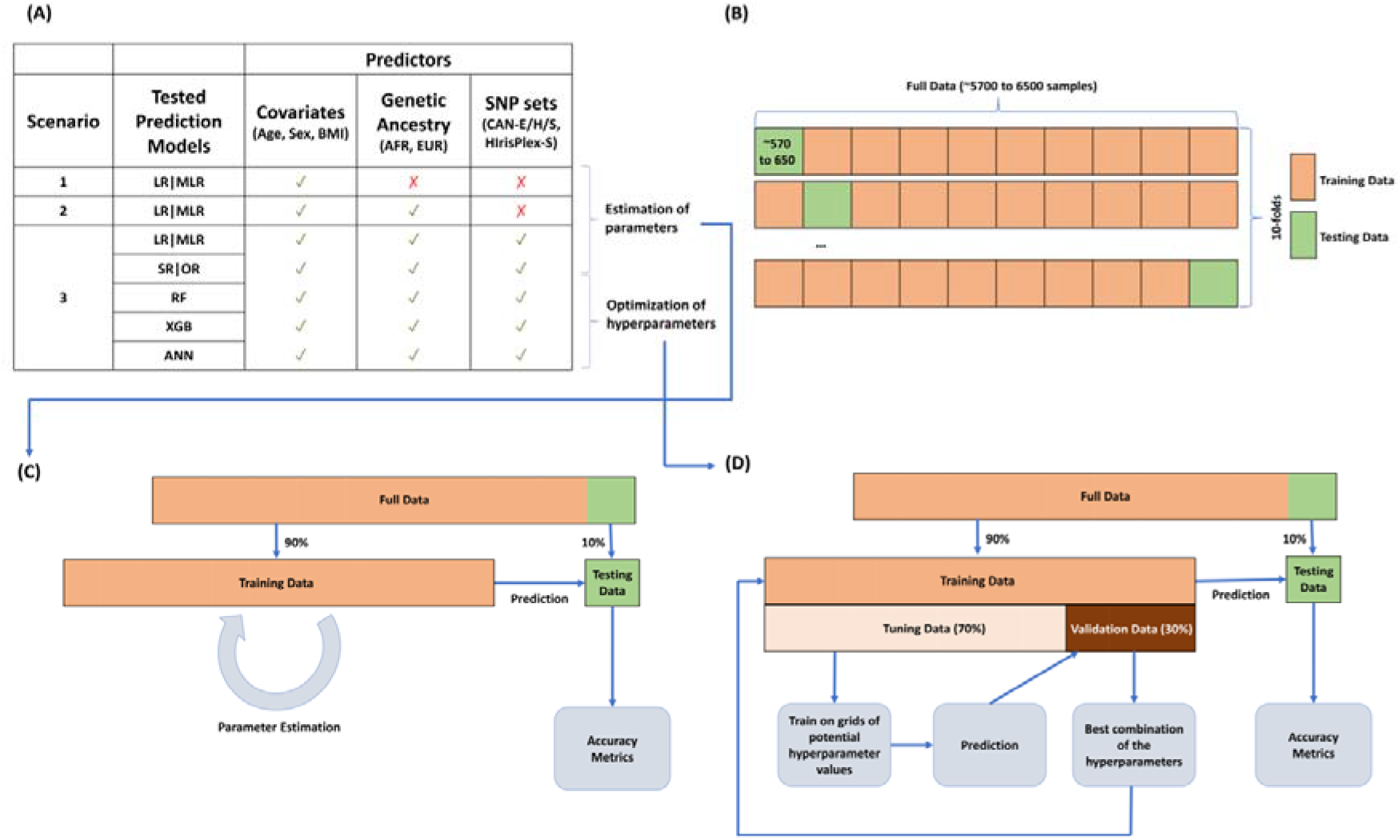
Study overview. **(A)** Models tested, predictors used and prediction methods: multinomial regression (MLR), linear regression (LR), ordinal regression (OR), stepwise regression (SR), random forests (RF), extreme gradient boosting (XGB) and artificial neural network (ANN). **(B)** 10-fold cross-validation: the full data is randomly split into 10 equally-sized data sub-groups. For each of the 10 sub-groups, the estimation of model parameters **(C)** or optimization of model hyperparameters **(D)** was performed on a pool of the nine remaining sub-groups.

For the third (full) model, in addition to regression, the following statistical and machine-learning methods were used, in order to evaluate their relative performance for prediction: Random Forest (RF), Extreme Gradient Boosting (XGB), Artificial Neural Network (ANN), Ordinal regression (OR) and Stepwise regression (SR). We provide more details on their implementation in Supplementary method S2.

### Evaluation of prediction accuracy

To measure prediction accuracy in quantitative traits, we used the coefficient of determination (*R*^2^, the proportion of phenotype variance that is explained by the model, measured as 1 – *SS_res_/SS_tot_*, where *SS_res_* and *SS_tot_* respectively stand for the residual and total sum of squares). For categorical traits, we used a metric denoted “accuracy” that is the proportion of correctly classified individuals in confusion matrices. For these traits, we also computed the Area Under the ROC Curve (AUC – ranging from 0.5 to 1) as well as the expected accuracies from two benchmark strategies (either using the categories’ frequency – PropStrat, or always guessing the most frequent category – maxP; see Supplementary Method S3) for comparison.

Accuracy of prediction was evaluated using 10-fold cross validation (10-fold CV) (Fig. 1B). For this, the full dataset is split into 10 approximately equal subsets based on a stratified sampling on the trait (so that trait distribution is similar across subsets). Each of the 1/10^th^ subsets is used as test data for evaluating prediction accuracy of methods trained using the other 9/10^th^ of the data. For regression methods (MLR, LR, SR, OR), coefficient parameters are estimated in the training data (Fig. 1C). For Machine Learning models, the training data is further split into a tuning data (70% of the training data) and validation data (30% of the training data). The parameter space for these methods are tuned creating a grid of all possible combinations of the hyperparameters and the particular combination producing best result on the validation data is selected as the set of optimal combination for the whole training data (see Supplementary Method S2). All possible permutations of binding nine folds for training produces 10 different train-test data combinations and the hyperparameters are tuned based on that subsequently gives rise to 10 prediction accuracy results (Fig. 1D). These measures of goodness of fit are used in a boxplot, or the average of them is used as a single prediction accuracy metric of the method. This helps us in avoiding inflation in the results and the predictions are more robust to small changes in the data.

### Prediction of MI in Native American individuals of unknown phenotype

Genotype and geo-localization data for 104 Native American individuals (with an estimated >99% Native American ancestry) from 16 Native American populations were available from a previous study[40]. We analysed genotypes for these ‘pure’ Native American individuals, predicted their MI, and regressed these predicted values on the amount of solar radiation at the site of population sampling. We trained two RF models (one with CAN-S and one with HIrisPlex-S) using 550 CANDELA individuals with >= 80% native ancestry and using sex as the only covariate. Solar radiation levels were defined as insolation incident on a horizontal surface (in kWh/m^2^/day) as reported in the NASA Surface meteorology and Solar Energy (SSE) Web site (https://eosweb.larc.nasa.gov/sse/) (data previously used in[31]).

## Results

Below we compare the performance of HIrisplex-S with SNP sets devised here for the prediction of eye (CAN-E), hair (CAN-H) and skin (CAN-S) from summary statistics of a pigmentation GWAS performed in the CANDELA sample (Materials and Methods and Supplementary method S1). The sets devised here consist of: 56 (CAN-E), 101 (CAN-H) and 120 (CAN-S) pigmentation-associated SNPs and are detailed in Supplementary table S2. Consistent with the genetic correlation of eye, hair and skin pigmentation, some SNPs are shared across the CAN-E/H/S sets, as well as with the HIrisPlex-S set. The overlap between these four SNP sets is shown in Figure 2.

**Figure 2.**
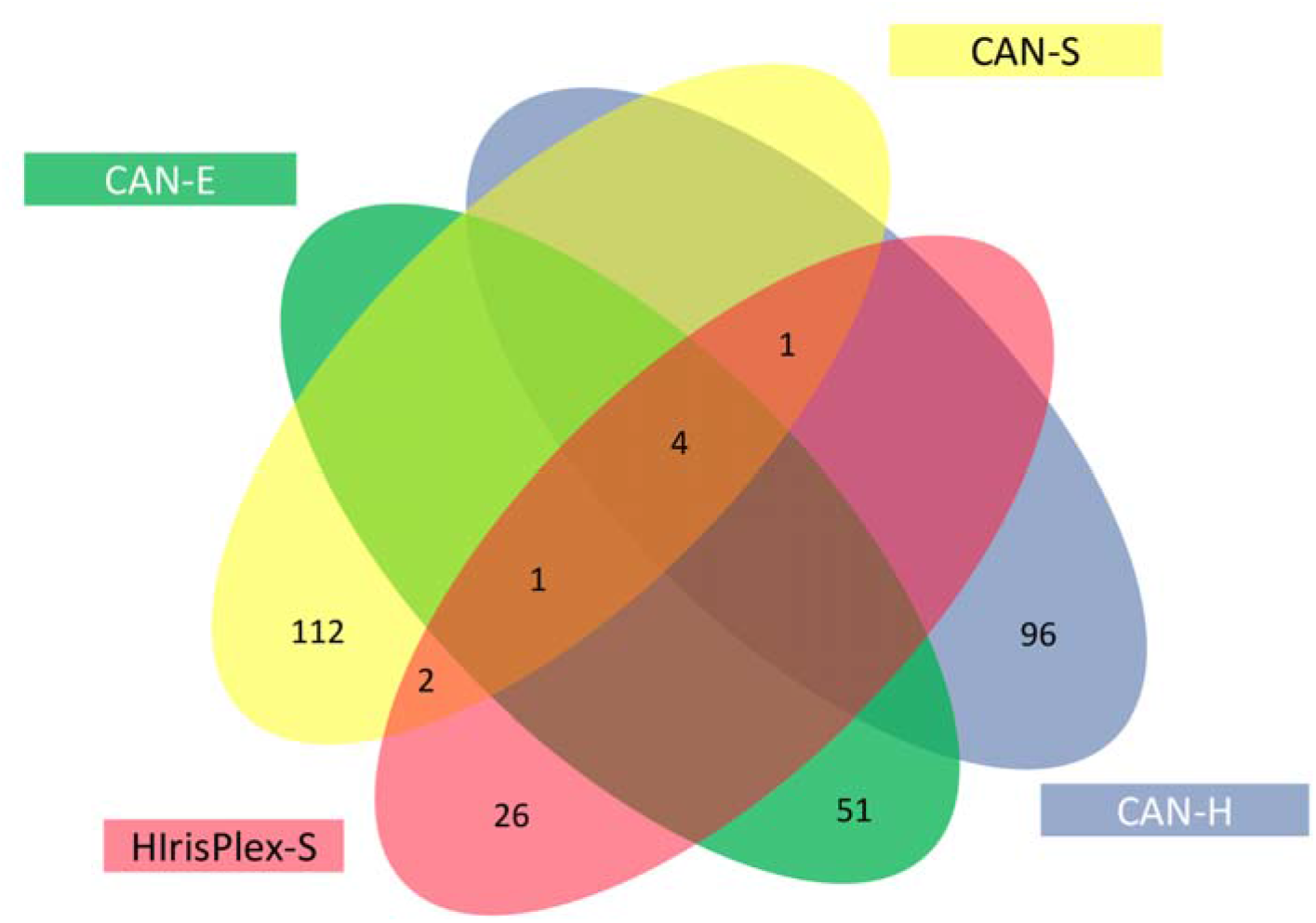
Overlap between SNP sets used for prediction of pigmentation traits. CAN-E, CAN-S and CAN-H refer, respectively, to SNP sets designed here for the prediction of eye, skin and hair pigmentation, based on a GWAS performed in the CANDELA sample[31]. HIrisPlex-S is a SNP assay developed for simultaneous Eye, Hair and Skin color prediction[14]. Numbers refer to SNPs shared between SNP sets.

### Prediction Accuracy in relation to models, methods and pigmentation SNP sets

Figure 3 presents the accuracy of prediction for various phenotypes of eye, hair and skin color. For categorical traits, the baseline model (i.e. including only non-genetic predictors: age, sex and BMI), reaches 84.9% and 81.7% accuracy (proportion of correctly classified individuals) for eye and hair color, respectively. That level of accuracy is actually also reached by always guessing the phenotype to be the most frequent category (maxP strategy, see Supplementary table S3 and Supplementary Method S2). This high accuracy obtained by a deterministic strategy probably relates to the highly skewed trait distribution in the CANDELA individuals: ~ 82% having black/dark brown hair and 85% having Brown/Black eyes. Alternately, randomly guessing the phenotypes based solely on the frequency of the traits (PropStrat; cyan line in Figure 3) also yields good levels of accuracy (~74% and ~69% for categorical eye and hair color, respectively).

**Figure 3.**
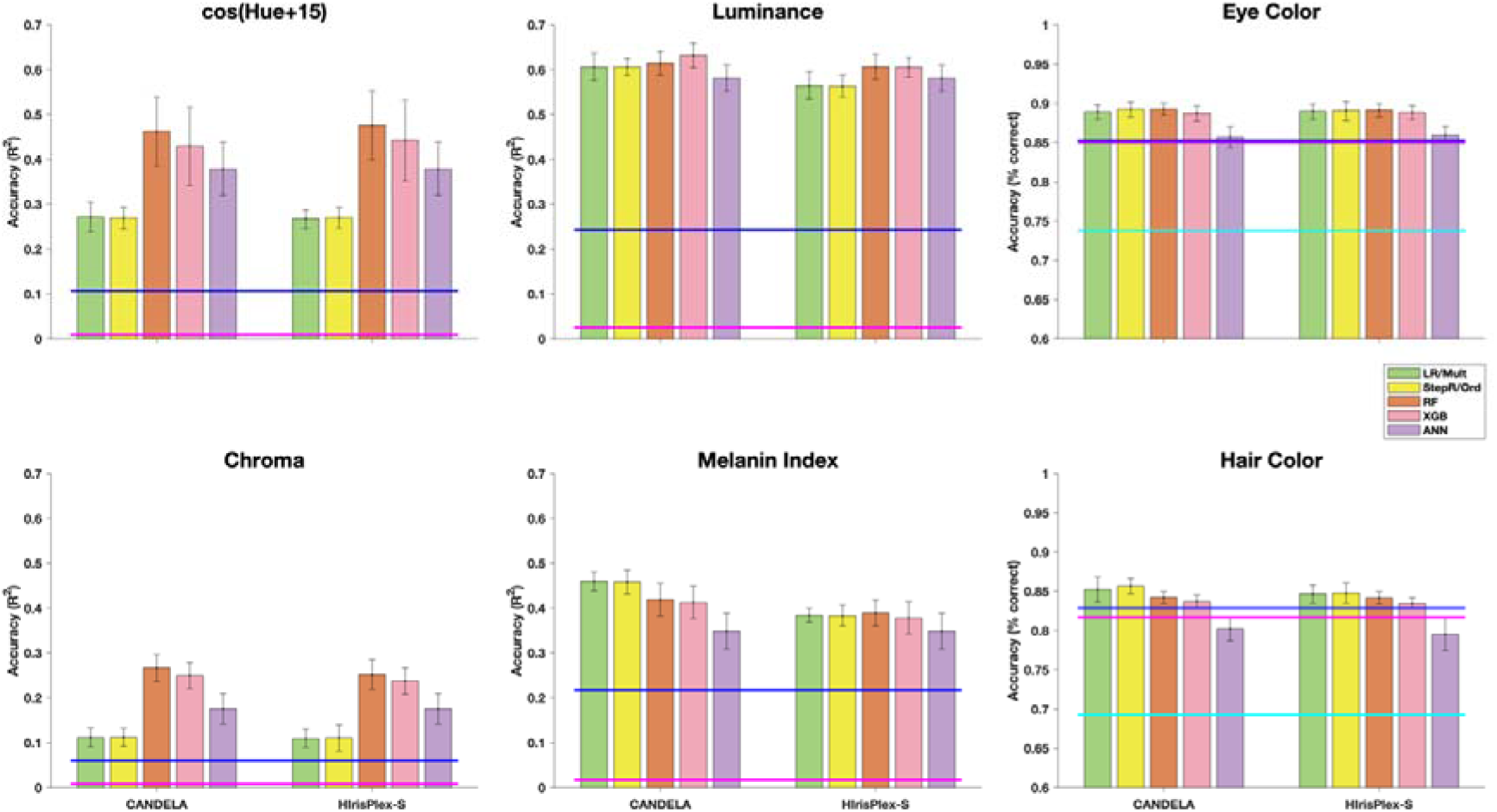
Prediction accuracy in relation to models, methods and pigmentation SNP sets. For continuous traits (Hue (transformed), Luminance, Chroma and Melanin Index; top and middle panels) we used R^2^ as measure of prediction accuracy. For categorical traits (Eye and Hair color; bottom panels) accuracy is the proportion of correctly classified individuals. Magenta and blue lines indicate the accuracy obtained when only non-genetic predictors or non-genetic + genomic ancestry are included in regression models, respectively. For categorical traits, the performance of a random guessing strategy (PropStrat) was also evaluated (cyan line). For these traits, the average accuracy of the deterministic maxP strategy is numerically the same as the accuracy obtained when only non-genetic predictors are used (magenta line), hence is not shown separately in this figure. For the full prediction model (non-genetic predictors + genetic ancestry + pigmentation SNPs) the performance of regression and four additional prediction methods was evaluated (bars are colored: green = LR/MLR; yellow = SR/OR; brown = RF; pink = XGB; purple = ANN). Detailed numerical values are given in Supplementary Table S3. The pigmentation SNP set incorporated in the prediction models is indicated at the bottom of the plots.

It is important to keep the maxP strategy in context when assessing prediction performance for categorical traits, since it represents how skewed the trait distribution is – a binary trait with a frequency distribution of 90% and 10% of the two categories will have 90% accuracy under the simplest maxP strategy (even though its sensitivity will be 0 for the rare category; see Supplementary Method S2). Thus, a skew in the trait distribution causes an upward shift in accuracy of prediction methods, especially those methods which are biased to the most frequent category, making them appear better-performing than they actually are. The PropStrat strategy is comparatively less biased as it gives proportional weight to the rare category (hence non-zero sensitivity for this category), and thus has lower accuracy than the maxP strategy. It is therefore a better benchmark to compare the performance of other strategies for assessing their gain in accuracy. Conversely, a comparison of those strategies to the maxP benchmark better represents their relative change in incorrect classification rather than the relative gain in correct classification (i.e. gain in accuracy).

Although the accuracies of these basic strategies are already high due to our skewed trait distributions, adding genetic ancestry to the model has a further impact, especially for hair color: it decreases the proportion of error by ~6% (from 18.3% of error to 17.1% - detailed numbers in Supplementary table S3). Then the further addition of pigmentation SNPs has an even larger effect: the remaining proportion of errors decreases by another ~16% and ~27% respectively for hair and eye color. It is also noticeable that the gain in prediction brought by SNPs relatively to that brought by genetic ancestries is much larger for eye color (~14*x*) than for hair color (~3*x*).

For continuous traits, we observe a large increase in prediction accuracy (R^2^) when genetic ancestry is incorporated in regression models, relative to the baseline (including only non-genetic predictors). Furthermore, when pigmentation SNP sets are incorporated in the regression model, accuracy usually more than doubles over that obtained with genetic ancestry plus non-genetic covariates (green bar versus blue line in Figure 3). When using this full prediction model, lowest LR prediction accuracy was observed for Chroma (R^2^ ~0.12) and highest for Luminance (R^2^ ~0.58), two quantitative estimates of eye color variation.

Comparing different prediction methods for the full model (i.e. incorporating all predictors) we do not observe large differences in performance, with the exception of a relatively lower accuracy of regression for Hue and Chroma. For those two traits, RF markedly outperforms regression methods, more than doubling the accuracy of LR in the case of Chroma. For the two other continuous traits (Luminance and Melanin Index), regression models are as effective as machine learning models (Figure 3). We also note that those rank similarly throughout categorical and continuous traits: RF is almost always better than the other tree-based model (extreme gradient boosting; XGB) and artificial neural networks (ANN) always underperform compared to tree-based models.

Regarding the SNP sets tested, we observe little difference in prediction accuracy across traits and methods although the number of SNPs is larger in the sets that we designed (56 to 120 SNPs) than in HIrisPlex-S (34 SNPs). Continuous skin pigmentation (Melanin Index) stands nonetheless as an exception: for that specific trait, CAN-S consistently outperforms HIrisPlex-S (particularly with regression methods). We have further assessed the impact of limiting the number of SNPs from the CAN sets to the top 10, 20 or 30 SNPs. A reduction in the number of SNPs had little effect for most traits, to the noticeable exception of Melanin Index for which considering the top 30 SNPs (i.e. 25% of the CAN-S SNP set) yields less than 90% of the prediction R^2^ achieved with the whole set (see Supplementary table S7). Also, we ranked the SNPs by their variable importance (returned by the RF models) for each trait (see Supplementary table S4 and figure S3). Three SNPs, from three known pigmentation genes (SLC45A2, HERC2 and SLC24A5), are consistently among the top ones throughout traits and are common to HIrisPlex-S and the 3 CAN SNP sets.

### Prediction accuracy at varying levels of European/Native American ancestry

Since a substantial fraction of individuals in the CANDELA sample have minimal African ancestry, we sought to evaluate prediction accuracy specifically for varying levels of European/Native American ancestry in the CANDELA sample. To this aim, we pooled individuals with negligible African ancestry in ~20% ancestry bins (so that the smallest pool size included at least 570 individuals) and examined prediction accuracy in the pools (see Supplementary table S5). Furthermore, for categorical traits, we ensured that at least two trait categories, each with >20 individuals, were observed in the pools, which led to withdraw the most Native-American pool. We assessed prediction accuracy using RF, a full model (i.e. equation 3 but without genetic ancestry as predictor) including only pigmentation SNPs having >1% MAF in the pool of individuals being tested.

For the categorical traits, there is a drop in prediction accuracy (from ~95% to ~70%) at increasing European ancestry (Figure 4). However, as European ancestry increases there is greater accuracy relative to random guessing based on trait frequency, probably reflecting the trait being less variable at higher Native American ancestry levels. For the quantitative eye pigmentation variables (particularly H and L, Figure 4), as the percentage of European ancestry increases there is a trend for an increase in trait variation (red line in Figure 4) and also in prediction R^2^. For skin pigmentation (MI) we observe an opposite trend in trait variability in relation to ancestry, relative to hair/eye color: variation in MI decreases at increasing European ancestry. There is also a trend towards an increase in the performance of the CAN-S SNP set at decreasing European ancestry: in individuals with <20% European ancestry CAN-S has an accuracy that is nearly twice that observed for HIrisPlex-S. Although CAN-S tends to outperform HIrisPlex-S in most comparisons, it is only for MI that such a large difference in performance was observed. In summary, across all pigmentation traits we observe a gain in prediction accuracy for the Native American/European ancestry bins showing greater phenotypic diversity, with CAN SNPs markedly outperforming HIrisPlex-S only for MI in individuals with low (<20%) European ancestry.

**Figure 4.**
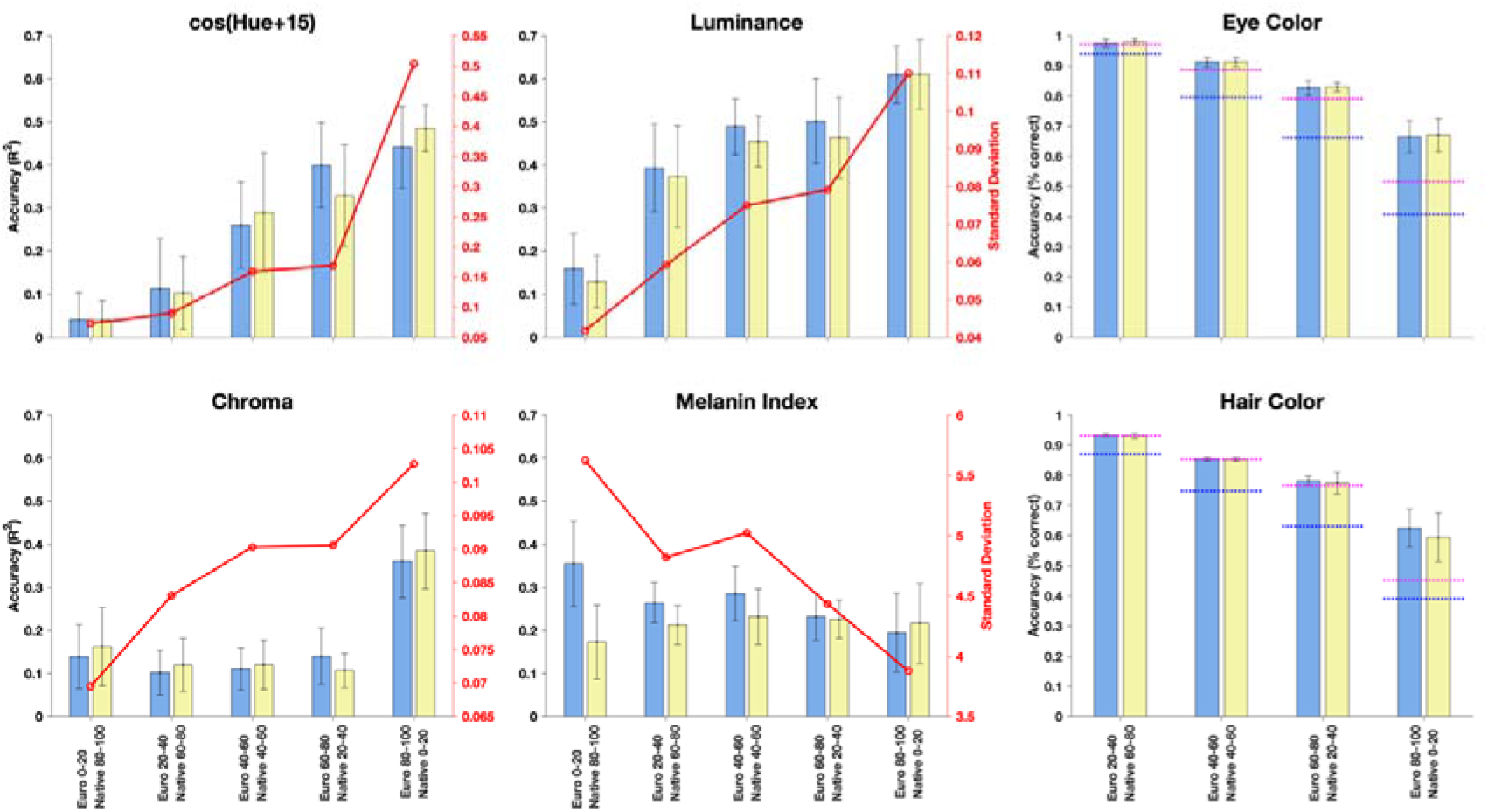
Prediction accuracy for individuals with varying Native American/European admixture. Prediction was assessed in individual bins varying ~20% in admixture (bottom axis; for eye and hair pigmentation <20 individuals with >80% Native Ancestry were available in each trait category, thus preventing estimation of prediction accuracy). Colored bars indicate accuracy obtained with Random Forest models using non-genetic + pigmentation SNP sets as predictors (R^2^ being used as prediction measure for quantitative traits: Hue, Luminance, Chroma and Melanin index). Blue bars indicate the CAN-E/H/S SNP sets. Yellow bars indicate the HIrisPlex-S set. The standard deviation of the quantitative traits in each ancestry bin is indicated as a red line. For the categorical traits (Eye and Hair Color), accuracy (proportion of correctly classified individuals) is used as the metric for prediction measures. Accuracy obtained without genetic predictors using a guessing strategy is indicated with a horizontal blue line for Proportional Strategy (random guessing) and magenta line for maxP (deterministic guessing). Detailed numerical values are given in Supplementary Table S8 and S9.

### Prediction accuracy in CANDELA relative to other population samples

Several studies have examined accuracy of prediction of the HIrisPlexS SNP set for categorical pigmentation traits using an online tool (https://hirisplex.erasmusmc.nl, referred to here as HIrisPlexS-Online), which implements MLR prediction models trained in a reference dataset (consisting mostly of Europeans)[7,11,14]. Table 1 compares the published estimates with those we obtained here with the CAN and HIrisPlex-S SNP sets using our implementation of MLR models trained in the CANDELA data (for skin color, we transformed MI into a 3-level categorical variable: fair, intermediate and dark; as described in Materials and Methods). For a more accurate comparison to HIrisPlexS-Online, these models were trained using SNPs only, without basic covariates or ancestry.

**Table 1.** Overall Accuracy (%) and trait AUC (%) for categorical eye (A), hair (B) and skin (C) color obtained here and in other studies (% frequency of trait in sample is shown in parentheses).

| Sample | CANDELA<br>N=6,529 |  |  |  | Venezuela/Brazil<sup>b</sup><br>N = 99 |  | Europeans<sup>d</sup><br>N=2,364 |  |
| --- | --- | --- | --- | --- | --- | --- | --- | --- |
| SNP set |  | CAN-E<sup>a</sup> | HIrisPlex-S | HIrisPlexS-Online |  |  |  |  |
| Overall Accuracy |  | 89 | 89 | 87 |  |  |  | 83 |
| 1. Blue/gray | (3) | 93 | 93 | 93 | (12) | 85 | (68) | 91 |
| 2. Intermediate | (13) | 89 | 89 | 85 | (23) | NA | (10) | 73 |
| 3. Brown/black | (85) | 92 | 92 | 90 | (64) | 91 | (23) | 93 |

(B)
| Sample | CANDELA<br>N=6,495 |  |  |  | Europeans<sup>c</sup><br>N = 385 |  |
| --- | --- | --- | --- | --- | --- | --- |
| SNP set |  | CAN-H | HIrisPlex-S | HIrisPlexS-Online |  |  |
| Overall Accuracy |  | 85 | 84 | 56 |  | 71 |
| 1.Red | (0) | NA | NA | NA | (25) | 90 |
| 2. Blond | (3) | 94 | 92 | 90 | (54) | 75 |
| 3. Brown | (16) | 82 | 81 | 65 | (9) | 72 |
| 4. Black | (82) | 87 | 85 | 80 | (12) | 78 |

(C)
| Sample | CANDELA<br>N=6,526 |  |  |  | World-wide<br>N=1,159<sup>e</sup> |  |
| --- | --- | --- | --- | --- | --- | --- |
| SNP set |  | CAN-S | HIrisPlex-S | HIrisPlex-S-Online |  |  |
| Overall Accuracy |  | 73 | 72 | 26 |  | 87 |
| 1. Fair | (38) | 83 | 81 | 78 | (92) | 97 |
| 2. Intermediate | (59) | 77 | 77 | 32 | (3) | 83 |
| 3. Dark | (3) | 86 | 84 | 81 | (5) | 96 |

HIrisPlexS-Online: https://hirisplex.erasmusmc.nl
<sup>a</sup> Obtained using a model with only SNPs as predictors (no covariates) and with MLR as prediction method.
<sup>b</sup> Obtained using HIrisPlex as described in Freire-Aradas et al. 2014[43].
<sup>c</sup> Obtained using HIrisPlex-S as described in Branicki et al 2011[44], here accuracy is calculated from a 4-way classification setup.
<sup>d</sup> Obtained from the specificity reported in Liu et al 2009[10]
<sup>e</sup> This prediction values reported in Walsh et al 2017[13] were obtained from the prediction of the 194 samples from Maronas et al 2014[42]

Prediction accuracy estimates for eye color have been reported for HIrisPlex-S-Online in a Latin American sample (99 individuals from Venezuela and Brazil) and in a European sample[10,14,15]. The light (blue/gray) and dark (Brown/Black) color categories have similar prediction accuracies across studies (~90-93%), except for the prediction of light eye color reported for HIrisPlex-S-Online in the Venezuelan/Brazilian sample, where accuracy is much lower (85%). The main difference in eye-color prediction across studies lies in the intermediate category. No intermediate eye colors were predicted by HIrisPlex-S-Online in the Venezuelan/Brazilian sample. Our predictions in the CANDELA sample have markedly higher accuracy than reported for HIrisPlex-S-Online in Europeans, both when the prediction model was trained in the CANDELA data or in the reference HIrisPlex-S data (respectively 89% and 85%, versus 73% in Europeans). Prediction accuracy, in the CANDELA sample, of the CAN-S and HIrisPlex-S SNP sets was identical.

Estimates for hair color prediction accuracy using HIrisPlex-S-Online have been reported for a European sample[44]. Prediction accuracy estimates obtained here for the CANDELA sample are higher than reported in Europeans for all hair colors, except in the case of intermediate hair-color (i.e. brown) predicted with HIrisPlex-S-Online. The highest hair-color prediction accuracy was consistently obtained with the CAN-H SNP set, although the difference relative to the HIrisPlex-S, with models trained in the CANDELA data, is small.

Concerning categorical skin color, we observe that predictions from HIrisPlex-S-Online have markedly lower accuracy in the CANDELA sample than reported for a worldwide sample. Model training in the CANDELA dataset increases prediction accuracy substantially both for HIrisPlex-S and CAN-S, although the accuracy values obtained are still below the published estimates for HIrisPlex-S. As for hair color, the CAN-S SNP set consistently outperforms HIrisPlex-S, but only marginally.

### Portability of models for pigmentation prediction in individuals with high Native Ancestry

Considering the impact of training datasets in the performance of HIrisPlex-S (Table 1), we specifically examined the portability of RF models developed in two training datasets with extreme differences in ancestry (extracted from the CANDELA sample – see Supplementary figure S4): (i) a highly European training dataset (European ancestry >= 80% and Native American ancestry < 20%) and (ii) a highly Native training dataset (European ancestry < 20% and Native American ancestry >= 80%). We examined the performance of the resulting prediction models in a subset of the highly Native test dataset (Figure 5) in a cross-validation scheme. We observe that models developed in the highly Native training dataset have a better performance than those developed in the highly European training dataset for Chrome, Luminance and MI. The most striking difference in performance is seen for MI, where the model trained with highly Native data has a prediction accuracy ~6 times that of the model trained in highly European data (Figure 5). Hue is the only trait for which the model trained in the highly European dataset outperforms the model trained in the highly Native dataset, but prediction accuracy in this case is extremely low (<2%).

**Figure 5.**
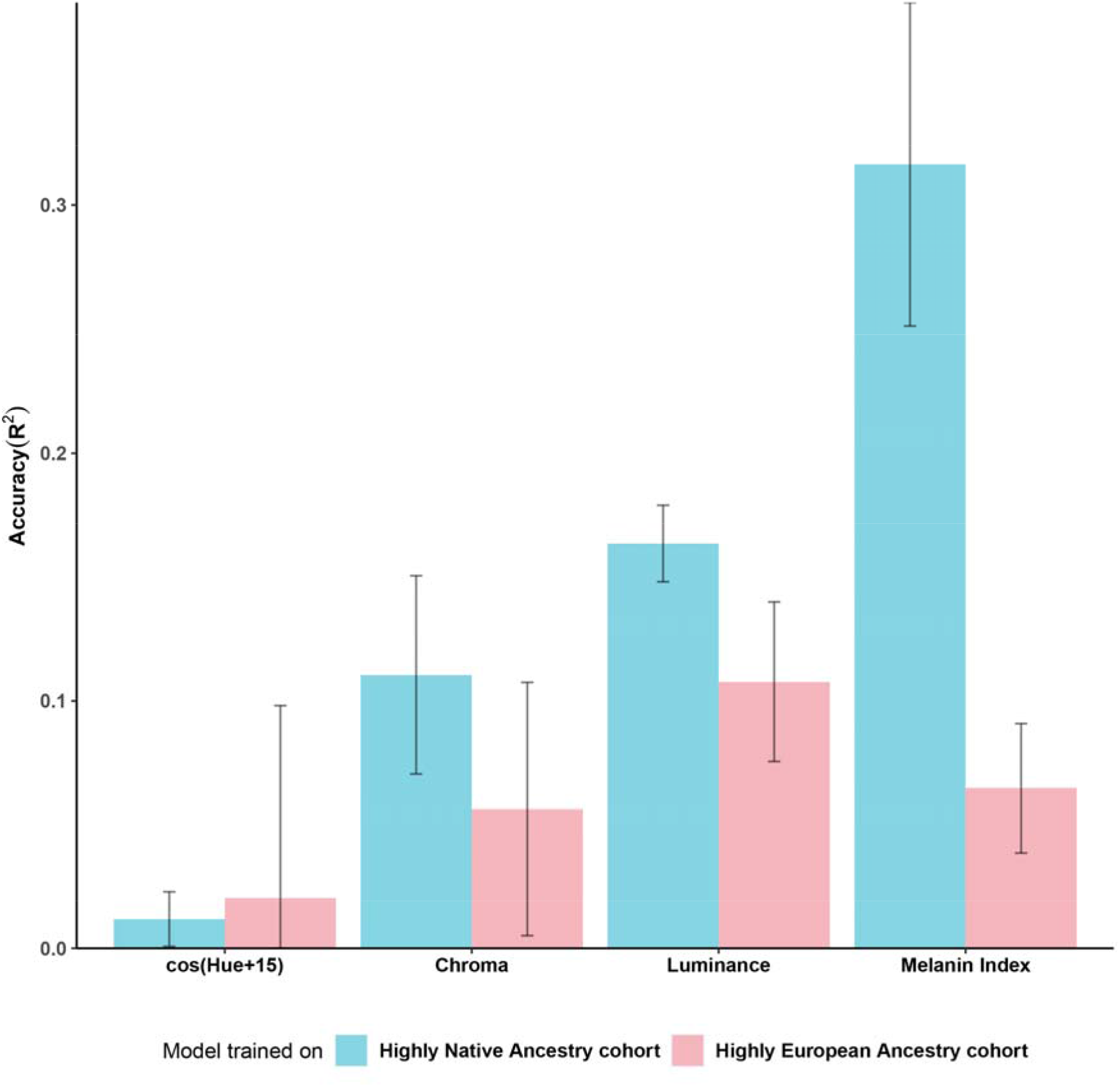
Portability of prediction models trained in highly European/Native American cohorts. For each continuous trait (cos(H+15), C, L and Melanin Index) we compare prediction accuracy on the same test data (a highly Native ancestry cohort). The prediction models were trained either on a highly Native (Blue) or highly European (Pink) cohort established from the CANDELA sample. For testing, we created equally-sized 4-folds for each pool of individuals. We built the RF models using three of the folds and evaluated prediction accuracy in the left-out fold from the Native ancestry cohort (see Supplementary figure S4).

### Prediction of skin pigmentation in Native Americans

We examined prediction performance of the CAN-S and HIrisPlex-S sets in a highly Native dataset independent of CANDELA by predicting MI in 117 individuals from 17 Native American populations[40]. As above, we trained RF prediction models using CANDELA individuals with >= 80% native ancestry. Since performance could not be measured directly in this dataset (due to the lack of phenotypic data), we examined the correlation of predicted skin pigmentation (MI) with solar radiation levels at the site of population sampling (Figure 6). Previous surveys of skin pigmentation in native populations from across the world have found a correlation between skin pigmentation and solar radiation[51], an observation that has been interpreted as the result of selection throughout human evolution. In the Native Americans examined here we obtained correlations of 0.543 (p-value 2 x 10^−9^) and 0.163 (p-value 0.1) with the CAN-S and HIrisPlex-S SNP sets, respectively (Figure 6).

**Figure 6.**
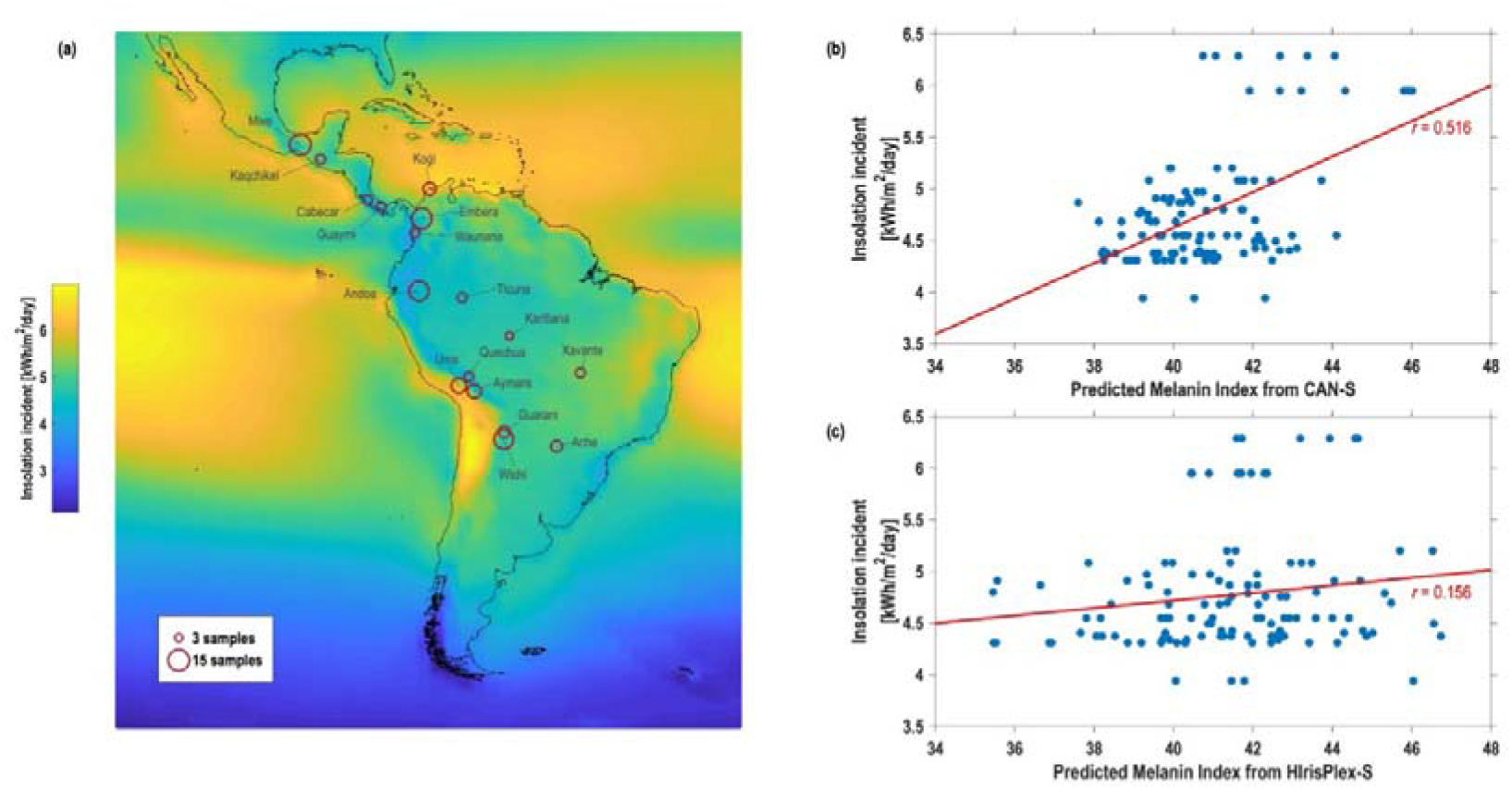
Solar radiation levels and skin pigmentation (MI) predicted with CAN-S and HIrisplex-S SNP sets. **(a)** Annual average of insolation incident on a horizontal surface (kWh/m^2/day - data from NASA Surface meteorology and Solar Energy, 2008) and location of the Native American population sampled. The predicted MI and solar radiation levels at the sampling site for 104 individuals from 16 Native American populations is shown in **(b)** CAN-S and **(c)** HIrisPlex-S.

## Discussion

Our results agree with previous studies in finding that regression and RF generally outperform other prediction methods (MLR being the approach implemented in HIrisPlex-S-Online). Owing to their tree-based structures, RF and XGB implicitly model the underlying genetic interaction across SNPs, and studies have noted the presence of epistasis (SNP-SNP interactions) between major pigmentation SNPs to some degree for various pigmentation traits, suggesting that prediction accuracy can be further increased by specifically allowing for interaction between SNPs[23,24,52]. Since tree-based models implicitly model epistasis including complex higher-order interactions, the difference of their prediction accuracy to additive linear/logistic models may also shed light on the genetic architecture of the traits. It is tempting to hypothesize that SNP interactions may have a more substantial impact on the genetic architecture of Hue and Chroma phenotypes[31,53], especially considering that these traits are relatively more non-linear in nature (e.g. Hue is an angle, i.e. a circular trait)[52]. Furthermore, the differences in prediction accuracy are much larger from linear to tree-based models (up to +0.19 gain in R^2^ – see Supplementary table S3) than they are from HIrisPlex-S to CAN-E (up to +0.014 gain in R^2^) whereas these two SNP sets only have 5 SNPs in common. We might therefore expect that such relevant interactions would be limited to a handful of SNPs, consistent with the literature which primarily observed interactions between major pigmentation SNPs[23,24,31]. Conversely, we may expect a greater additive polygenicity[23,25] in the case of skin pigmentation. Linear models overperform tree-based models for that trait (Figure 3) and the number of SNPs in the model has an impact on the prediction accuracy, as suggests the +0.07 gain in R^2^ from HIrisPlex-S (34 SNPs) to CAN-S (120 SNPs). In line with these observations, we also noted that prediction accuracies are relatively lower for Melanin Index than for other continuous traits when constraining the number of SNPs in the model (see Supplementary table S6 and S7).

Previous studies[31,36,54,55] have shown that genetic ancestry correlates with pigmentation, probably resulting from the variable allele frequency across continents of certain pigmentation-related alleles. Consistent with this, we observe that continental genetic ancestry has significant predictive power (Figure 3, Supplementary table S3) and that the SNPs retained for prediction rank among the most correlated to continental ancestry components (see Supplementary table S2). Inclusion in the prediction models of pigmentation SNP sets selected from the GWAS further increases predictive power, markedly for quantitative pigmentation traits. Candidate SNPs on an average add twice the prediction accuracy of that brought in by the genetic ancestry for the quantitative traits. For the categorical traits, the difference is smaller but comparatively candidate SNPs decreases the proportion of incorrect classification by a larger amount than does continental genetic ancestry, especially for eye color.

The difference in predictive power for categorical traits is partly the result of their intrinsic lower informativeness relative to quantitative traits. This is particularly so for hair and eye color, as the CANDELA sample has a highly skewed phenotypic distribution for these traits: 82% of individuals in the CANDELA sample are assigned to the darkest category for eye and 85% for hair color. This skewed phenotypic distribution is consistent with the fact that world-wide lightly pigmented eyes and hair are essentially Western Eurasian traits, while there is considerable variation in skin pigmentation outside Europe[23–28]. That is, the occurrence of lightly pigmented hair and eyes in the CANDELA sample is essentially a reflection of its partly European ancestry. This may explain why the HIrisPlex-S SNP set built primarily in Europeans performs equally well to the CAN SNP set built on our more diverse admixed samples, and matches what has been observed in the literature for predicting eye and hair color in non-European populations[23,32,33].

By contrast, prediction of skin pigmentation is influenced by the genetic architecture of this trait in all three populations contributing to admixture in Latin America (Native Americans, Europeans and Africans). Consistent with this, our analysis along a Native American-European ancestry gradient show the highest gain in prediction accuracy for the ancestry bins with the greatest phenotypic diversity: that is, the highest European bin for eye/hair color and the highest Native American bin for skin color. In the bin with lowest European ancestry, there is hardly any gain in prediction accuracy for eye/hair color over the deterministic maxP strategy, as almost all individuals in that bin are in the highly pigmented category. On the other hand, this bin has the highest variation in skin pigmentation, also shows the greatest accuracy for the CAN-S, and this SNP set strongly outperforms HIrisPlex-S prediction for this ancestry bin (Figure 4) – all these facts together emphasize that skin color prediction in admixed Latin Americans is substantially influenced by the genetic architecture of this trait in non-European populations. Our observation of a stronger correlation of predicted skin with solar radiation levels in pigmentation in Native Americans for the CAN-S set, relative to HIrisPlex-S is also consistent with this, and with literature reporting comparatively poor portability of European-based pigmentation prediction models on non-European populations[23,32,33].

Although quantitative traits are intrinsically more informative than categorical ones, anthropological and forensic applications of pigmentation prediction are usually interested in a few discrete categories, often just two (e.g. blue v. non-blue eyes) or three (light, intermediate or dark color). In that setting we find that for the CANDELA dataset there is little difference in the prediction performance of HIrisPlex-S (which, in addition, was shrunken from 41 SNPs to 34 polymorphic SNPs in our dataset), relative to the CAN SNP sets developed here. However, we observe a strong impact of the training dataset used for optimizing the prediction methods, both in the extreme scenario explored in Figure 5, and when comparing the prediction accuracies we obtained here with those published previously (Table 1). These observations question the portability across human populations of models developed using a training dataset with a different genetic makeup. This is most sharply illustrated by the difference in performance between the online implementation of HIrisPlex-S and the results we obtained when training models with this SNP set in the CANDELA data, particularly for skin pigmentation categories (Table 1). These observations thus caution against the use of the online implementation of HIrisPlex-S for the prediction of pigmentation traits in non-Europeans.

## Acknowledgments

We thank the volunteers for their enthusiastic support for this research. We also thank Alvaro Alvarado, Mónica Ballesteros Romero, Ricardo Cebrecos, Miguel Ángel Contreras Sieck, Francisco de Ávila Becerril, Joyce De la Piedra, María Teresa Del Solar, William Flores, Martha Granados Riveros, Rosilene Paim, Ricardo Gunski, Ana Angélica Leal Barbosa, Sergeant João Felisberto Menezes Cavalheiro, Major Eugênio Correa de Souza Junior, Wendy Hart, Ilich Jafet Moreno, Paola León-Mimila, Francisco Quispealaya, Diana Rogel Diaz, Ruth Rojas, and Vanessa Sarabia, for assistance with volunteer recruitment, sample processing and data entry. We are very grateful to the institutions that allowed the use of their facilities for the assessment of volunteers, including: Escuela Nacional de Antropología e Historia and Universidad Nacional Autónoma de México (México); Universidade Federal do Rio Grande do Sul (Brazil); 13° Companhia de Comunicações Mecanizada do Exército Brasileiro (Brazil); Pontificia Universidad Católica del Perú, Universidad de Lima and Universidad Nacional Mayor de San Marcos (Perú). We also acknowledge the Centre de Calcul Intensif d’Aix-Marseille for granting access to high-performance computing resources. We thank Manfred Kayser for assistance with Online HIrisPlex-S.

## Data availability

Summary statistics from the GWAS analyses on which were established the three CAN SNP sets used in this study were previously deposited at GWAS central (http://www.gwascentral.org/study/HGVST3308).

