## Supplementary Material for "Prediction of Eye, Hair and Skin Color in Admixed Populations of Latin America"

**SUPPLEMENTARY MATERIALS**

### Supplementary Methods

#### Supplementary method S1. Details on the construction of CAN-E, CAN-S and CAN-H SNP sets

We have established three SNP sets for prediction of one categorical (3-levels eye color) and three continuous (Hue, Chroma, Luminance) traits of eye color (CAN-E), of one continuous trait of skin pigmentation (melanin index – CAN-S) and of one categorical trait of hair color (3-levels hair color – CAN-H). The constitution of each one of these SNP sets was based on the genome-wide association results of the corresponding trait previously published in Adhikari et al. (2019). For the specific case of CAN-E, Luminance was retained as the reference traits among the four studied traits. Here below we detail the selection procedure. This procedure was independently applied for each of the three sets.

**Pre-selection of SNPs.** All SNPs suggestively associated to trait $y$ in the discovery GWAS was pre-selected (*p*-value <1E-5). In this discovery GWAS, the SNP-phenotype association was independently tested for about 9M SNPs with a linear model involving 8 covariates (sex, age and 6 genetic PCs).

**Computation of the predictive power of each SNP taken independently**. For the $i^{th}$ pre-selected SNP, we computed its predictive power $\rho_{i}^{2}$ as the proportion of phenotypic variance explained by that SNP, thus $\rho_{i}^{2}=\left( corr\left( \varepsilon_{y},\varepsilon_{g} \right) \right)^{2}$, where $\varepsilon_{y}$ and $\varepsilon_{g}$ are the residuals of the linear regressions of the GWAS covariates ($X\equiv$ sex, age and 6 genetic PCs) respectively on the phenotype ($y$) and on the genotype of the $i^{th}$ SNP ($g_{i}\equiv$ the number of counts of the minor allele of each individual $j$, i.e. $g_{i,j}=\left\{ 0,1,2 \right\}$).

**Clustering of pre-selected SNPs in linkage disequilibrium (LD).** Most often, a signal of association gathers various suggestive and/or significant SNPs from the same LD block. We clustered these SNPs into clumps using PLINK’s function --clump with default parameters. This algorithm builds clumps of variants around an index SNP (i.e. the most significant SNP of a certain region) by gathering any variant that is either located max. 250 kb from the index SNP or has a LD R2 with that index SNP greater or equal than 0.50. The number of clumps obtained ($N$) were: 701 (CAN-E), 1,471 (CAN-S) and 207 (CAN-H).

**Selection of one SNP per clump.** We retained the SNP with highest predictive power within each clump.

**Ranking of SNPs on their conditional predictive power.** The SNP with highest predictive power is ranked first. Then, we iteratively compute the conditional predictive power of any SNP that is not yet ranked. This conditional predictive power is the proportion of phenotypic variance explained by a given (unranked) SNP accounting for the usual covariates ($X$) and the genotypes of the SNPs that are already ranked at this stage ($M^{k})$. In a more formal way, we apply the following iterative ranking to all SNPs but the one with highest predictive power.

$$for k=2 \to N \left\{ \right.$$

$M^{k}\equiv$ a matrix containing the genotypes of the $k-1$ SNPs already ranked

$v^{k}\equiv$ a vector containing the predictive power of the$N-k+1$ unranked SNPs, conditioned on the $k-1$ SNPs already ranked

$$for i=k \to N \left\{ \right.$$

$g_{i}\equiv$ the genotype of the *i*^th^ unranked SNP

$\hat{\beta_{k}^{y}}\equiv$ the estimates of the linear regression of $\left( \begin{matrix} X & M^{k} \end{matrix} \right)$ on $y$

$\hat{\beta_{k}^{g_{i}}}\equiv$ the estimates of the linear regression of $\left( \begin{matrix} X & M^{k} \end{matrix} \right)$ on $g_{i}$

$$\varepsilon_{y}=y-\left( \begin{matrix} X & M^{k} \end{matrix} \right)\cdot\hat{\beta_{k}^{y}}$$

$$\varepsilon_{g}=g_{i}-\left( \begin{matrix} X & M^{k} \end{matrix} \right)\cdot\hat{\beta_{k}^{g_{i}}}$$

$$v_{i}^{k}=\rho_{i}^{2}=\left( corr\left( \varepsilon_{y},\varepsilon_{g} \right) \right)^{2}$$

$$\left. \right\}$$

$$j=argmax(v^{k})$$

The *j*^th^ unranked SNP is ranked at rank *k*

$$\left. \right\}$$

**Determination of the set of SNPs with highest prediction accuracy.** We retain the set $\Omega_{i}$ of SNPs ordered by rank of conditional predictive power ($\Omega_{i}=\left\{ {SNP}_{1},\cdots,{SNP}_{i} \right\}$) that has the highest prediction accuracy. Hence, the search for the optimal subset among selected SNPs is limited to a total of $N$ subsets including SNPs by decreasing predictive power.

To that end, for each set $\Omega_{i}$, we compute the prediction accuracy (R^2^) of a Scenario 3 (see section *Prediction scenarios tested and approaches used, equation 3*) using RF and following the cross-validation scheme of our study (see Figure 1: the whole data is split in 10 folds and a model is trained on each leave-one-out). Before retaining the set $\Omega_{i}$ that has the highest prediction accuracy, the vector of prediction accuracies is locally averaged with a smoothing kernel in order to avoid potential local maxima. We eventually retain the set $\Omega_{i}$ for which $i$ is minimal among those whose ${R2}_{i}\geq\max\left( R2 \right)-0.001$(see Supplementary Figure S5).

For the particular case of CAN-H, the trait is categorical (3-levels hair color). The optimal set $\Omega_{i}$ was thus determined for each of the three 1-versus-2 category and the largest one was eventually retained.

#### Supplementary method S2. Statistical and Machine learning models used for prediction

**Multinomial Logistic Regression (MLR)** is the generalization of the binary logistic regression to a multiclass problem. Where the probability of an individual belonging to a certain class is

$\Pr(Y_{i}=1)= \frac{exp(b_{1}.X_{i})}{1+ \sum_{k=1}^{K-1} exp(b_{k}.X_{i})}$,

$\Pr(Y_{i}=2)= \frac{exp(b_{2}.X_{i})}{1+ \sum_{k=1}^{K-1} exp(b_{k}.X_{i})}$,

…,

$\Pr(Y_{i}=(K-1))= \frac{exp(b_{K-1}.X_{i})}{1+ \sum_{k=1}^{K-1} exp(b_{k}.X_{i})}$, and

$\Pr\left( Y_{i}=K \right)=1-\Pr\left( Y_{i}=1 \right)-\Pr\left( Y_{i}=2 \right)-\ldots-Pr \left( Y_{i}=(K-1 \right))$,

$\boldsymbol{b}_{\boldsymbol{k}}$ is the set of the regression coefficients associated with outcome k, is the set of independent variables and is the categorical phenotype of the individual i. The regression coefficients are estimated using an iterative algorithm and the final category predictions are derived using modal probabilities from MLR. In our analysis, k takes 3 values for both the categorical phenotypes - Blue/gray, Intermediate and Brown/black levels for eye color and blond, dark blond/light brown and brown/black for hair color and the independent variables are the basic covariates, estimated continental genetic ancestries (European and African) and SNPs from different SNP sets.

**Linear Regression (LR)** is a linear method to model the relationship of a quantitative dependent variable on a set of independent variables. A straight line is fitted across the datapoints that minimizes the error sum of squares and gives us the equation for prediction given the values of the independent variables. For the continuous traits (H, L, C, Melanin Index) we used LR for Scenario 0, 1 and 2 where the variables used each Scenario are already explained earlier.

**Random Forest (RF)** is an ensemble of multiple decision trees, where the modal voting is chosen as the prediction for a categorical variable or a mean of the predictions from each of the trees for a continuous variable. Biggest advantage of RF is that this method corrects for the tendency of decision trees to overfit using the hold out samples/ out of bag samples in the training process of the individual trees. For our prediction models, we used the R package ‘randomForest’ and tuned the trees using a grid search over ntree and mtry parameters. ‘ntree’ determines how many trees should be there in the forest whereas ‘mtry’ determines how many variables should be sampled before a new split. We took a parameter set (50, 100, 150, … , 750) for ntree and (3 to 40) for mtry. The best combination of ntree and mtry is selected from the validation data and applied on the test data to arrive at the accuracy metric. RF was also used for the selection of SNPs to be included in the CANDELA prediction set.

**Extreme Gradient Boosting (XGBoost)** is a sequential tree ensemble method where each new tree tries to explain the residual from the previous tree. The first tree of the series is modeled on the entire training data and each subsequent tree tries to make the model better. Thus this method can easily lead to overfitting. So we created a grid of hyperparameters where we took nrounds, early stopping rounds, max depth, eta, colsample by level, colsample by tree, subsample, min child weight and eval metric, alpha, gamma, lambda and ‘gbtree’ as booster to optimize the models. We took ‘nrounds’ values from 1 to 50 and ‘early stopping rounds’ values from (10, 15, 20) mandates the number of exact rounds the method will train if the accuracy does not improve on the validation set. Our objective was set as ‘reg:linear’ for the quantitative traits and ‘multi:softmax’ for categorical traits and evaluation metric was set as ‘rmse’ and ‘mlogloss’ respectively.

**Neural Network (NN)** is a circuit of neurons or directed graphs, that is composed of multiple layers of nodes, including one input and one output layer and a variety of hidden layers in between. Each node is connected to the previous or next or both layers through some edges, which carries a weight. The weights are optimized in a way to minimize the overall error rate of the network. We used 3 intermediate layers for all our models and for categorical traits the output layer had 3 neurons, one for each of the levels of both the traits (eye color and hair color) where for quantitative traits the output layer has only one neuron for the final prediction. The networks are compiled by a loss function and an optimization function. The loss functions were mean squared error for continuous traits and categorical cross-entropy for categorical traits and the optimization function was rmsprop. Activation functions also help us to shape the network. We used relu and tanh for continuous traits and additionally used softmax for categorical traits.

**Ordinal regression (OrdinalR)** is a type of regression analysis used for prediction of ordinal response variables. For ordinal regression we add the covariates and SNP information as our independent variables to the model where we assess the probability of each individual falling into one of the categories for eye color or hair color. Also the phenotypes are properly ordered to ensure there is a gradual change between the levels. Similar to MLR, an iterative procedure estimates the coefficients of the independent variables for the models and the final outcome is estimated as the category with highest probability estimated.

**Stepwise Regression (StepR)** is an iterative regression model which adds a new variable to the existing model or removes an existing variable from the model based on a prespecified criterion, which in this case is Akaike Information Criterion (AIC). This model gives us the precise set of variables needed to gain the optimum accuracy for the quantitative traits. At each step, this model checks for the best independent variable (from the set of remaining SNPs and covariates which are not already included in the model) to be added to the model only if it brings additional accuracy or removes any variable already added in the model that does not effectively increase the prediction.

#### Supplementary method S3. Benchmark Prediction Strategies

For the categorical variables, there are certain prediction strategies which are considered as benchmark for comparison of any newly proposed prediction strategy. The common two strategies are:

1. A deterministic strategy: always selecting the largest category by proportion of samples, here called maxP (Maximum Proportion).
2. A random guessing strategy: randomly assigning a category with probability equal to the proportions of each of the levels, here called PropStrat (Proportional strategy).

A discussion of their characteristics, including how they are affected by a skewed trait distribution, is provided in the methods. Here we provide some further details explaining those observations.

For the categorical variables, we create confusion tables which are the cross-tables of the predicted levels of the categorical dependent variables against the observed levels. The accuracy gives us the proportion of the correctly classified individuals from the confusion table.

Accuracy = (True Positives + True Negatives) × 100 / (Total number of observations).

But along with accuracy, sensitivity is another metric that helps us to understand the separate predictability of each of the levels of the categorical variable. We calculate Sensitivity for each level from the confusion table as the percentage of correctly classified individuals for that level.

Sensitivity = (True Positives) × 100 / (True Positives + False Negatives).

Though maxP always dominates over PropStrat in terms of accuracy (proved below), sensitivity for the non-majority categories is better for PropStrat. For maxP, the sensitivity for the non-majority groups always remains at 0 and for the majority group at 1. Where for PropStrat, the sensitivity for each group is the proportion of that category.

We were motivated to add these two strategies as benchmark in the case of categorical variables with skewed distributions, as it is very easy to reach a certain accuracy with a biased strategy like maxP without actually making any effort for predictive modeling, but at the cost of sensitivity. Therefore, to get a sense of additional accuracy gain with the prediction models using the covariates and genetic information over these benchmark strategies, the corresponding accuracy levels are indicated on all the accuracy plots of categorical traits, and their accuracy values are provided in the Supplementary Tables.

**Illustration with a binary trait:**

Let us consider a classification setup of a binary variable with levels A and B with n_1_ and n_2_ samples respectively. After prediction, say, we get m_1_ and m_2_ samples predicted as A and B respectively, out of which n_11_ and n_22_ are identified correctly. The confusion table would look like the following:

| Actual  Predicted | A | B | Row total (number of predicted samples) |
| --- | --- | --- | --- |
| A | n_11_ | n_12_ | m1 |
| B | n_21_ | n_22_ | m2 |
| Column total (number of samples) | n_1_ | n_2_ | Total = N |

The accuracy of the prediction approach would be (n_11_ + n_22_)/N, where total number of samples N = m_1_ + m_2_ = n_1_ + n_2_.

Sensitivity will be n_11_/n_1_ and n_22_/n_2_ for A and B respectively.

Accuracy for the two benchmark strategies will be maxP = max(n_1_/N, n_2_/N) and propStrat = (n_1_/N)^2^ + (n_2_/N)^2^.

Sensitivity of the two categories in the case of maxP is (n_1_/n_1_, 0/n_2_) if n_1_ > n_2_ or (0/n_1_, n_2_/n_2_) if n_1_ < n_2_, so (1,0) or (0,1) depending on n_1_ and n_2_, whereas for propStrat it is (n_1_/N, n_2_/N).

So for a very skewed distribution of the levels (where either n_1_/N or n_2_/N becomes very small and close to 0) then accuracy for maxP = max(n_1_/N, n_2_/N) also approaches 1. But for the minor level, the sensitivity remains to be 0. For propStrat accuracy also increases with a skewed distribution but it always remains less than that of maxP (proved below), whereas sensitivity always remains above zero.

**Accuracy comparison between maxP and propStrat**

Let us assume there are k levels of a categorical variable, where p_i_ denotes the proportion of the i-th level. So $\sum_{i=1}^{k} p_{i}=1$.

Now according to the Proportional Strategy (propStrat), the correct guesses for each of the groups are proportional to the weightage of that particular group, since labels are predicted randomly with probabilities equal to group proportions. Hence overall accuracy is:

propStrat = $\sum_{i=1}^{k} p_{i}^{2}$

≤ $\sum_{i=1}^{k} p_{i}.p_{max}, \mathrm{where} p_{max}= \max_{1\leq x\leq k} \left( p_{i} \right)$

= $p_{max} .\sum_{i=1}^{k} p_{i}$

= $p_{max} . 1$ = $\max_{1\leq x\leq k} \left( p_{i} \right)$ = maxP

So, we have proved that in terms of accuracy, maxP will always perform equal or better than propStrat. The equality holds when $p_{max}= p_{i}, for all i,1\leq i \leq k,$ that is when all the levels have same frequency and then maxP = propStrat = 1/k.

#### Supplementary method S4: Diversity Index

**Gini-Simpson Index:**

The diversity index (in Table S4) for categorical variables varies from 0 to 1, 0 meaning the data is completely alike and 1 meaning maximum variation within the categories where every observation belongs to a different category.

Gini-Simpson Index = 1 - $\sum_{i=1}^{R} p_{i}^{2}$, Where R is the number of levels of the variable and p_i_ is the sample proportion of level i.

Close observation reveals this diversity index is in fact the completement of the proportional strategy (propStrat) output, they adds up to 1 for each categorical variable. It also explains that higher the diversity, lesser chance of making a correct guess at random.

### Supplementary Figures

#### Supplementary figure S1. Frequency distribution of the quantitative traits

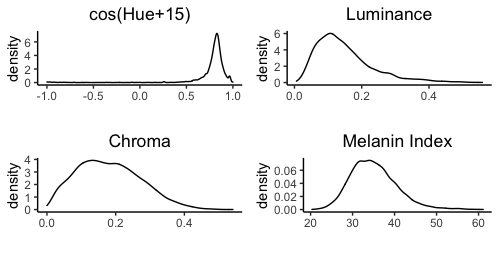

#### Supplementary figure S2: Melanin index thresholds defining categorical skin color

We created two ethnicity clusters using 100% European ancestry individuals (70 individuals) and individuals with African ancestry higher than native or European ancestry (23 individuals) with individuals having estimated genetic ancestry and profile pictures available and we manually ranked them in very pale, pale for the European cluster and dark and very dark from African cluster. Both clusters also included cases when the rater failed to classify any individual in any of these groups in particular. We plot the histograms of Melanin Index for all 4 groups of individuals (very pale, pale, dark and very dark). Finally, we identified the threshold of very pale and pale as our Melanin Index cut-off between fair and intermediate categories, also the lowest melanin index of the individuals in the dark category as the cut-off between intermediate and dark categories at 33 and 47 respectively.

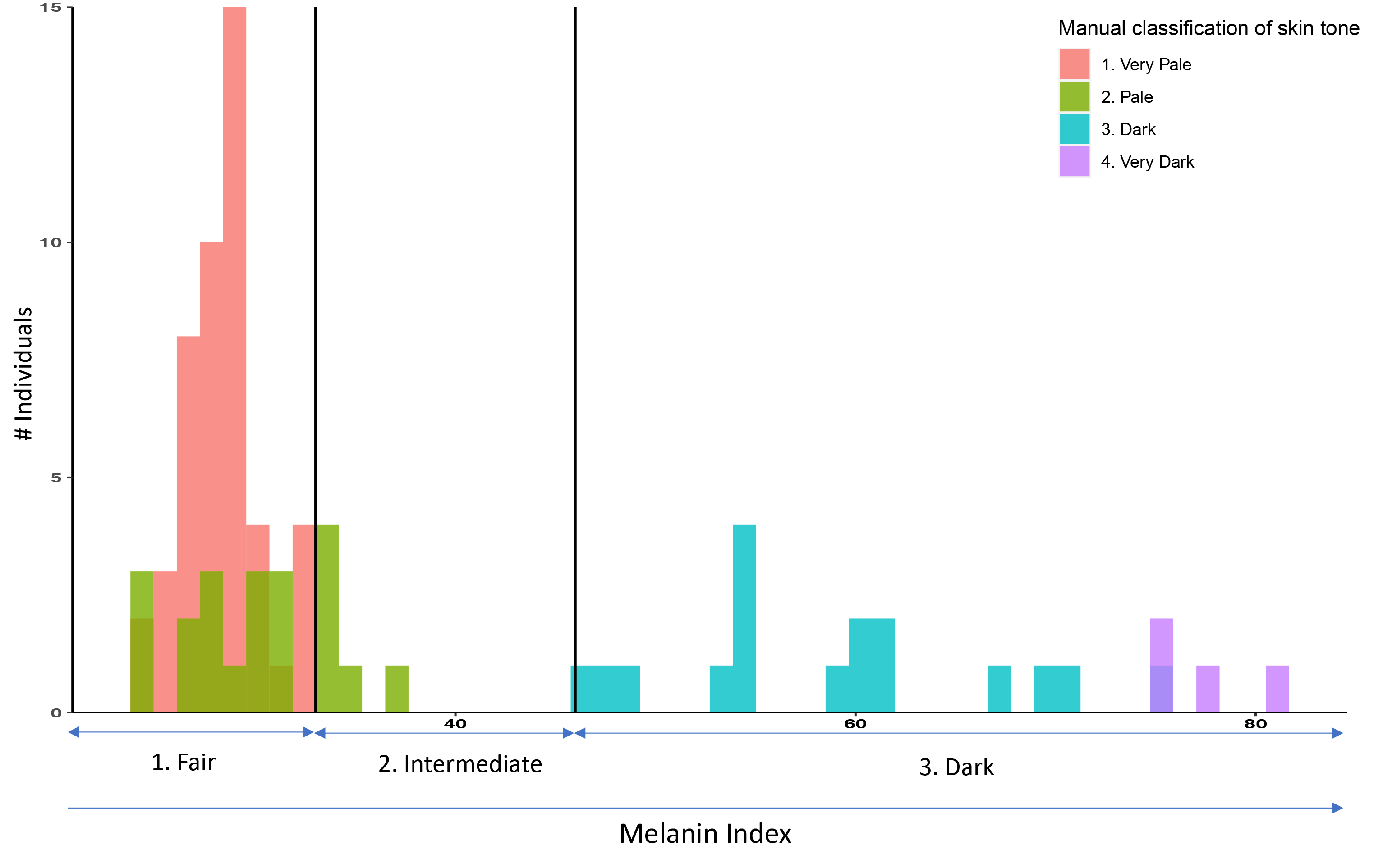

#### Supplementary figure S3. Variable importance and SNP sharing across sets.

We derived the variable importance from the random forest models in Equation 3 using CAN sets and retained the top 20 variables in each case (SNPs and covariates combined). The SNPs which are present also in the HIrisPlex-S assay has been denoted by green, SNPs present only in the CAN sets are denoted by red and covariates are highlighted in blue. We have used increase in mean squared error when a certain variable is dropped as our variable importance metric. The full list of CAN SNPs with their variable importance is provided in Supplementary table S2.

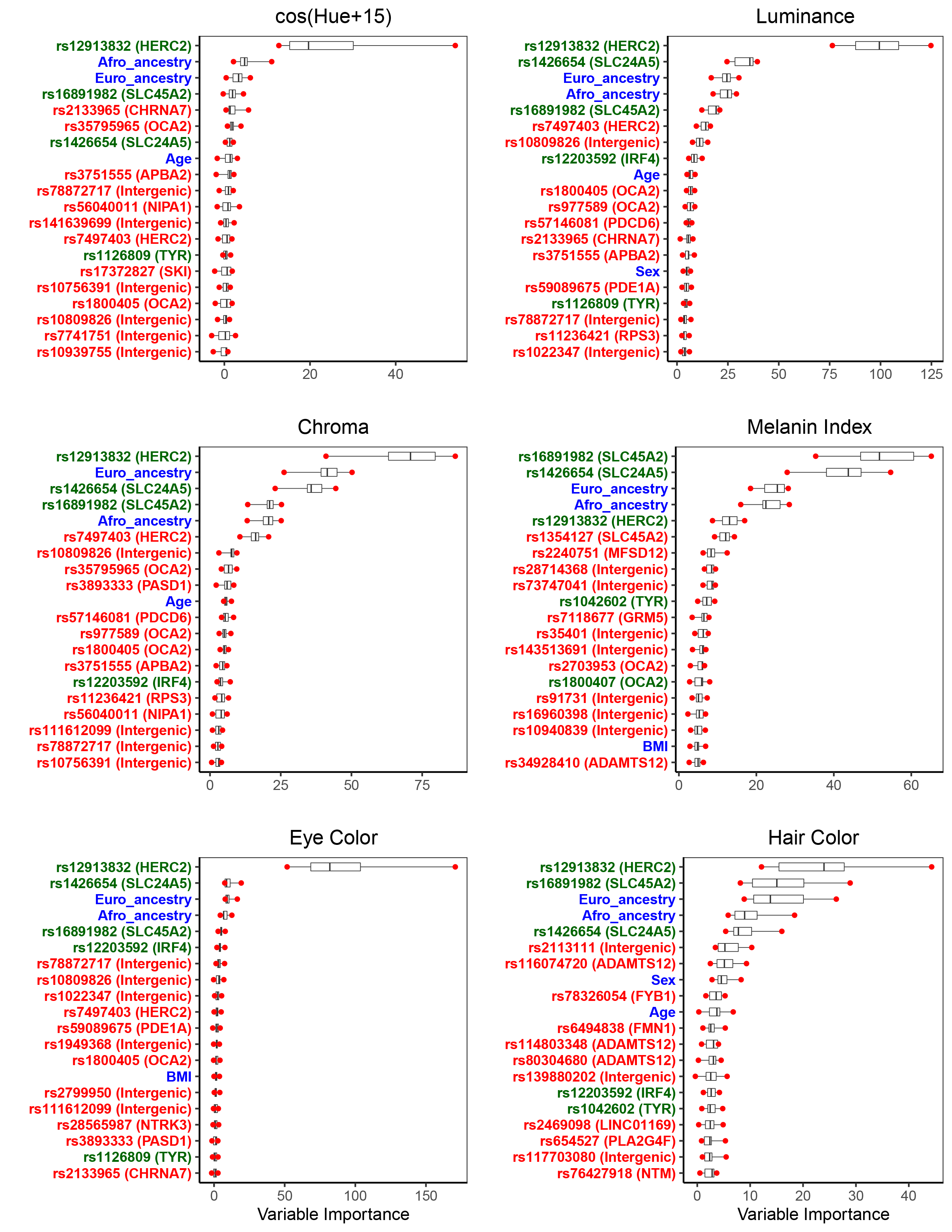

#### Supplementary figure S4. Modeling approach for ancestry portability.

The high European cohort data is trimmed to match the size of the high native cohort. Then both the datasets are divided in 4 equal folds, we train the random forest models on any 3 folds from one of the ancestry datasets in the training data and test the prediction accuracy on the remaining fold of the native ancestry data. So the models are trained on equal sized data from any of the extreme ancestry cohorts but the test data always remains to be highly native (for example, here fold 1 ,2 and 3 have been used to train the two models on the high native ancestry data and high European ancestry data – orange blocks and both the models are tested on the fold 4 – green bar of the high native ancestry data).

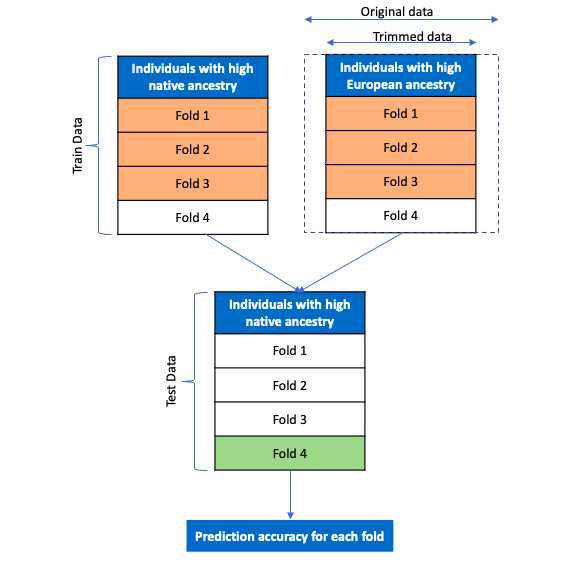

#### Supplementary figure S5. SNP selection to construct the CAN-E/H/S sets from SM1

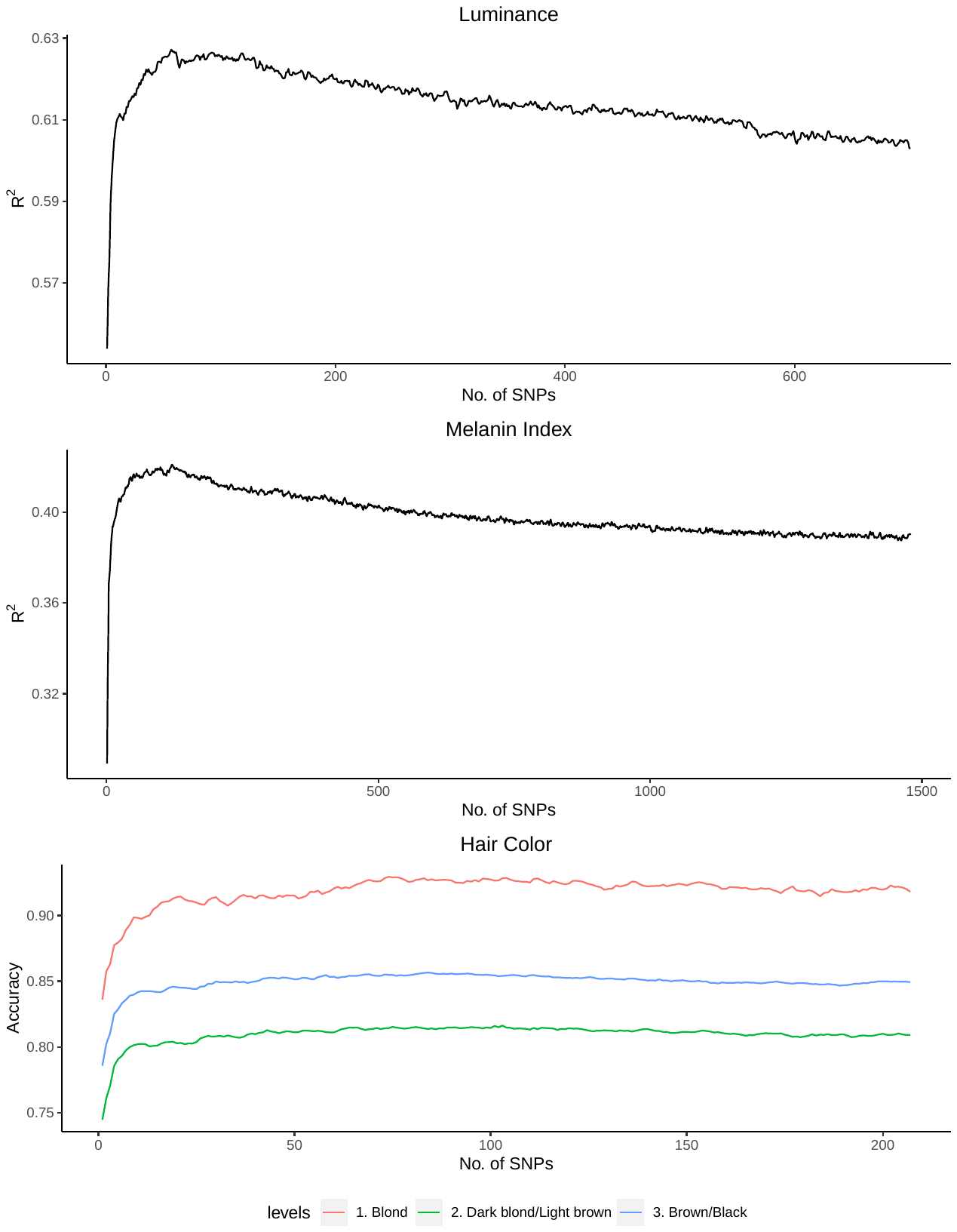

### Supplementary Tables

#### Supplementary table S1: List of all SNPs included in the HIrisPlex-S set

*See file “PREDICTION_PIGMENTATION_SupplementaryTables.xlsx”, tab “Table S1”*

#### Supplementary table S2: List of all SNPs included in the CAN-E/H/S SNP sets

*See file “PREDICTION_PIGMENTATION_SupplementaryTables.xlsx”, tab “Table S2”*

#### Supplementary table S3: Prediction accuracy numbers

*See file “PREDICTION_PIGMENTATION_SupplementaryTables.xlsx”, tab “Table S3”*

#### Supplementary table S4: Prediction accuracy from reduced models

*See file “PREDICTION_PIGMENTATION_SupplementaryTables.xlsx”, tab “Table S4”*

#### Supplementary table S5: Variable Importance in prediction for the covariates and CAN-E/H/S SNPs included in the pigmentation prediction models

*See file “PREDICTION_PIGMENTATION_SupplementaryTables.xlsx”, tab “Table S5”*

#### Supplementary table S6: Variable importance of the top 10, 20, 30 SNPs from Random Forest model for all 6 phenotypes

*See file “PREDICTION_PIGMENTATION_SupplementaryTables.xlsx”, tab “Table S6”*

#### Supplementary table S7: Ancestry pools distribution

*See file “PREDICTION_PIGMENTATION_SupplementaryTables.xlsx”, tab “Table S7”*

#### Supplementary table S8: Prediction accuracy for ancestry pools

*See file “PREDICTION_PIGMENTATION_SupplementaryTables.xlsx”, tab “Table S8”*

#### Supplementary table S9: Variability across ancestry pools

*See file “PREDICTION_PIGMENTATION_SupplementaryTables.xlsx”, tab “Table S9”*
